# TGFß/activin-dependent activation of Torso controls the timing of the metamorphic transition in the red flour beetle *Tribolium castaneum*

**DOI:** 10.1101/2023.08.04.551921

**Authors:** Sílvia Chafino, Roser Salvia, David Martín, Xavier Franch-Marro

## Abstract

Understanding the mechanisms governing body size attainment during animal development is of paramount importance in biology. In insects, a crucial phase in determining body size occurs at the larva-pupa transition, marking the end of the larval growth period. Central to this process is the attainment of the threshold size (TS), a critical developmental checkpoint that must be reached before the larva can undergo metamorphosis. However, the intricate molecular mechanisms by which the TS orchestrates this transition remain poor understood. In this study, we investigate the role of the interaction between the Torso and TGFß/activin signaling pathways in regulating metamorphic timing in the red flour beetle, *Tribolium castaneum*. Our results show that Torso signaling is required specifically during the last larval instar and that its activation is mediated not only by the prothoracicotropic hormone (Tc-Ptth) but also by Trunk (Tc-Trk), another ligand of the Tc-Torso receptor. Interestingly, we show that while Torso activation by Tc-Ptth determines the onset of metamorphosis, Tc-Trk promotes growth during the last larval stage. In addition, we found that the expression of *Tc-torso* correlates with the attainment of the TS and the decay of juvenile hormone (JH) levels, at the onset of the last larval instar. Notably, our data reveals that activation of TGFß/activin signaling pathway at the TS is responsible for repressing the JH synthesis and inducing *Tc-torso* expression, initiating metamorphosis. Altogether, these findings shed light on the pivotal involvement of the Ptth/Trunk/Torso and TGFß/activin signaling pathways as critical regulatory components orchestrating the TS-driven metamorphic initiation, offering valuable insights into the mechanisms underlying body size determination in insects.

## Introduction

How animals reach their final body size is a fundamental question in biology. Organism final body size depends on environmental cues as well as on the precise activation of genetic programs during development. In many animals, growth takes place mainly during the juvenile stage, and adult body size is therefore determined upon entering into adulthood [1]. Deciphering the molecular mechanisms underlying the timely decision to initiate adult maturation is, therefore, critical to understand how body size is controlled.

Hormones play an important role in the regulation of final body size, in part by coordinating the onset of adult maturation. For example, in holomeabolous insects, whose growth period is restricted to a series of larval molts that accommodate the increasing size of the body, the onset of metamorphosis is triggered by a sharp increase of the steroid hormone ecdysone upon reaching a size-dependent developmental checkpoint [2]. Ecdysone is synthesized in a specialized organ named the prothoracic gland (PG) through the sequential catalytic action of a series of enzymes encoded by the *Halloween* gene family. These include the Rieske-domain protein *neverland* (*nvd*) [3,4], the short-chain dehydrogenase/reductase *shroud* (*sro*) [5] and the P450 enzymes *spook* (*spo*), *spookier* (*spok*), *phantom* (*phm*), *disembodied* (*dib*) and *shadow* (*sad*) [6–11]. The expression of the Halloween genes is highly regulated, being up-regulated in the PG at the metamorphic transition by the integrated activity of several receptor tyrosine kinases (RTKs) [12]. These RTKs include Torso [13–15], Epidermal Growth Factor Receptor (Egfr) [16,17], Anaplastic Lymphoma Kinase (Alk) and PDGF and VEGF receptor-related (Pvr) [12], all acting through the Ras/Raf/Erk MAP kinase signal transduction cascade, and the Insulin receptor (InR) acting through the PI3K/Akt pathway [18–21]. Among these RTKs, Torso is of particular interest since it acts as a key transducer of critical environmental cues such as nutrition status and population density [15,22]. In order to exert its regulatory function in the PG, Torso binds the prothoracicotropic hormone (Ptth), a neuropeptide secreted from neuroendocrine cells located in the brain [13,23,24]. Although Ptth is considered the unique Torso ligand during postembryonic development, the fact that torso mutants showed a significant longer delay than Ptth null mutants [15], suggests the existence of additional ligands involved in the activation of Torso signaling. In fact, in addition to the Ptth, Torso possesses another ligand, Trunk (Trk), belonging to the cysteine knot growth factor superfamily, responsible for the activation of the pathway during the early embryo [14,25–27]. Although *trk* is not expressed during postembryonic stages of the fly *Drosophila melanogaster*, ectopic expression of a cleaved form of Trk in the PG is able to activate the pathway inducing precocious pupariation [26]. Nevertheless, to date, the activation of the Torso pathway by Trk has only been observed in the embryogenesis of all the studied species [25,26].

Despite the well-documented role of the Ptth/Torso pathway in triggering the onset of metamorphosis in *Drosophila* and *Bombyx mori*, the regulation of the pathway itself is less understood. Whereas Ptth synthesis and release from the neuroendocrine brain cells seems to depend on nutritional and environmental cues [13,18,20,28], the regulatory mechanisms that control the expression of *torso* is poorly understood. In this regard, it has been shown in *Drosophila* that the activity of the Transforming Growth Factor ß (TGFß)/Activin pathway in the PG is required for the proper expression of *torso* [29]. However, since depletion of TGFß/activin affects cell growth and morphology [29], it is plausible that *torso* regulation by this pathway is a collateral rather than a direct effect. In addition, in the beetle *Tribolium* and hemimetabolous insects such as *Gryllus* and *Blattella*, TGFß/activin has also been described as a negative regulator of the biosynthesis of the anti-metamorphic juvenile hormone (JH) [30–32], suggesting a possible link between the decay of JH at the end of larval development and the activation of the Torso signaling pathway at the onset of metamorphosis.

Here, we use the red flour beetle *Tribolium* to study the regulation of metamorphic timing by the Torso signaling pathway. During development, the size of *Tribolium* increases over several larval instars until reaching a critical size-assessment checkpoint - the *threshold size* (TS) - that instructs the larva to enter metamorphosis at the ensuing molt, thus ending the growth period [33]. In laboratory conditions, the TS is reached at the onset of the seventh larval instar (L7), and is associated with the decay of JH and the consequence down-regulation of the anti-metamorphic transcription factor *Krüppel-homolg 1* (*Tc-Kr-h1*). As a result, the stage-specific transcription factors *Ecdysone inducible protein 93F* (*Tc-E93*) becomes up-regulated triggering metamorphosis [33]. Unfortunately, although the described genetic changes that control the nature of the metamorphic transition have been studied in detail [33,34], the molecular mechanisms underlying the regulation of the metamorphic timing in *Tribolium* remain to be clearly defined.

In the present study, we uncover the role and regulation of *Tc-torso* in the control of the metamorphic timing in *Tribolium*. We show that Torso is required specifically in the last larval instar of the beetle for the synthesis of ecdysone that promotes the metamorphic transition. Remarkably, we found that during this period Torso signaling is activated not only by Tc-Ptth but also by Tc-Trk. However, whereas Tc-Torso activation by Tc-Ptth determines the onset of metamorphosis, Tc-Trk promotes growth during the last larval stage. Moreover, our experiments indicate that *Tc-torso* expression depends on larvae reaching the TS at the onset of the last larval instar, and that the sustained increased in the expression levels of *Tc-torso* during the last larval instar depends on the decay of JH titters. Interestingly, we found that the activity of Baboon (Tc-Babo) and Myostatin (Tc-Myo), two key components of the TGFß/activin signaling pathway, are required to repress JH synthesis and activate the TS-dependent induction of *Tc-torso* expression. Taken together, our results indicate that the Ptth/Trk/Torso and TGFß/activin pathways are critical component of the mechanism that controls the final body size of *Tribolium* by regulating growth and the timing of the metamorphic transition.

## Results

### Torso signaling regulates metamorphic timing in *Tribolium*

To investigate the role of Torso signaling in *Tribolium*, we first examined its temporal expression pattern during larval development. *Tc-torso* mRNA levels were low during early stages of larval development but strongly increased after entering into the last larval instar (L7), reaching the maximal level of expression at 24-48h, to decline thereafter until the end of the instar (Fig 1A). This result suggests that Torso signaling in *Tribolium* is necessary during the last larval stage of development. To test this possibility, we analysed the effects of blocking Torso signaling from early larval development by injecting dsRNA of *Tc-torso* in newly molted antepenultimate L5 larvae (*Tc-torso^RNAi^* animals). Specimens injected with dsMock were used as negative controls (*Control* animals). Under these conditions, *Tc-torso^RNAi^* larvae molted with proper timing to L6 and then to L7, but showed a significant developmental delay in the pupation time (Fig 1B). These results confirmed that Torso signaling in *Tribolium* is specifically required during the last larval instar to control the timing of the metamorphic transition.

**Figure 1.**
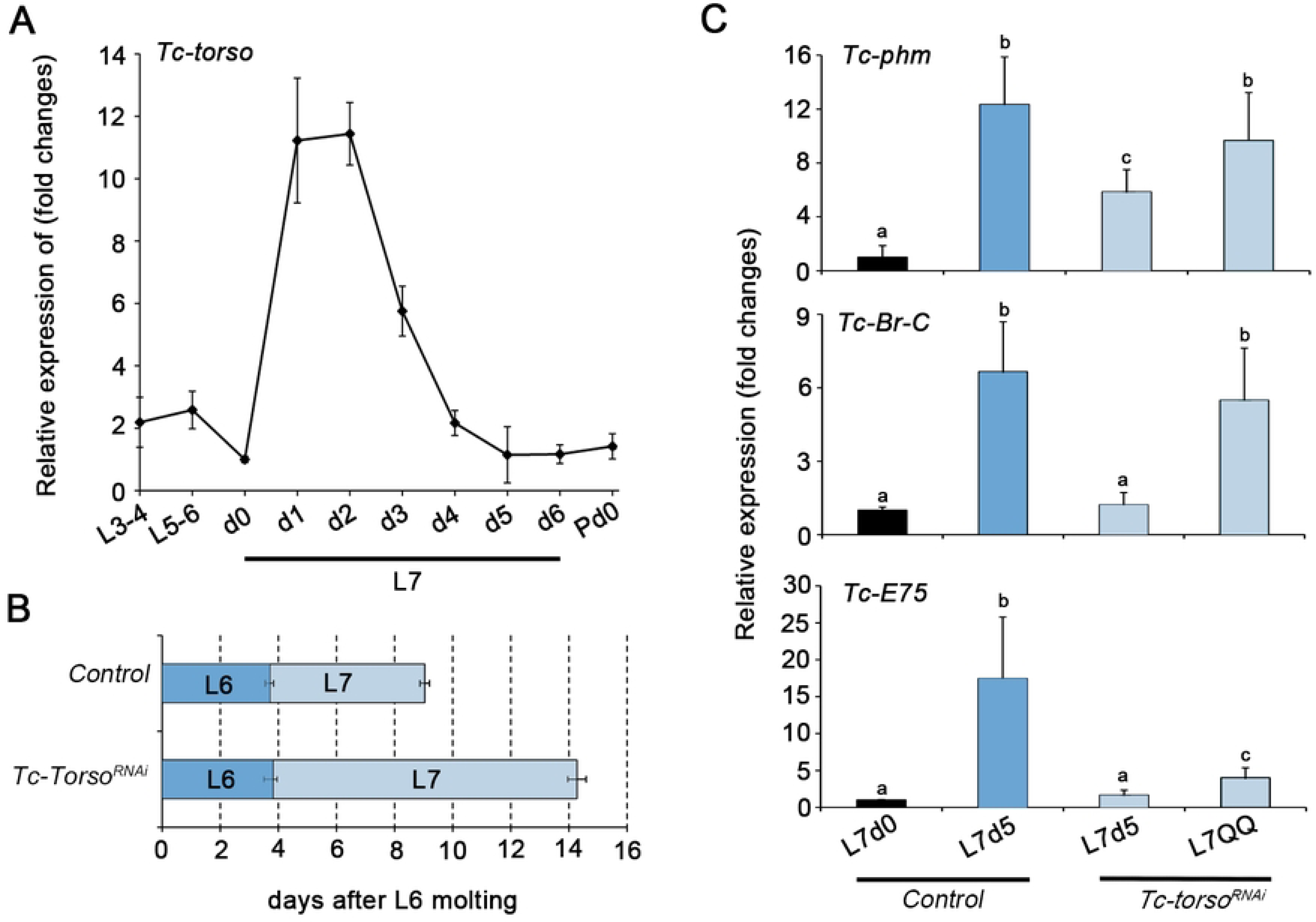
Tc-Torso is a critical regulator of pupariation in Tribolium. (A) Temporal changes in *Tc-torso* mRNA levels measured by qRT-PCR from L3-4 instar larvae to pupa. Transcript abundance values were normalized against the Tc-Rpl32 transcript. Error bars indicate the standard error of the mean (SEM) (n=5). (B) Developmental progression of newly molted L5 larvae after injection with either dsMock (*Control*) (n=30) or *dsTc-torso* (n=22). The bars represent the mean ± standard deviation (SD) for each developmental stage observed after the double-stranded RNA (dsRNA) injection. (C) Transcript levels of *Tc-phm*, *TcBr-C*, and *Tc-E75* measured by qRT-PCR in 0 and 5-day-old L7 Control larvae and 5-day-old L7 and quiescent (QQ) *Tc-torso^RNAi^* larvae. Transcript abundance values were normalized against the *Tc-Rpl32* transcript. Error bars indicate the SEM (n=9). Different letters represent groups with significant differences based on an ANOVA test (Tukey, p < 0.001).

To determine whether the pupation delay observed in *Tc-torso^RNAi^* larvae was caused by an ecdysone deficiency, we next analyzed the expression levels of the representative Halloween gene *Tc-phm*, as well as a number of well characterized ecdysone-dependent genes such as *Tc-Br-C* and *Tc-E75* that are used as proxies for ecdysone levels. Consistent with delayed pupation, *Tc-torso^RNAi^* larvae presented a significant delay in the expression of all the analyzed genes when compared to *Control* animals (Fig 1C). Altogether, these results indicated that Torso activation during the last larval stage of *Tribolium* is required to control the production of ecdysone that timely triggers pupa formation.

### Ptth and Trk activate Torso signaling during the last larval instar

To further characterize the function of Torso during the metamorphic transition, we knocked-down this receptor specifically in the last larval stage. Under this treatment, *Tc-torso*-depleted larva pupated with a delay of 7 days (Fig 2A). However, contrary to *Drosophila*, where depletion of either *Tc-Ptth* or *Tc-torso* resulted in bigger pupae [14], *Tc-torso^RNAi^* pupae were slightly smaller and lighter than *Control* pupae (Fig 2C-E). These results suggest the possibility that postembryonic activation of Tc-Torso (not only controls developmental timing but also growth rate) might not only depends on Tc-Ptth and the regulation of ecdysone synthesis but also in the direct control of growth rate. To further study this possibility, we analyzed the role of Ptth in Torso signaling activation in Tribolium. First, we measured the expression level of *Tc-Ptth* in last instar larvae, and found fluctuating levels with two peaks at day 2 and 4 (Fig 2C). Interestingly, depletion of *Tc-Ptth* in *Tribolium* (*Tc-Ptth^RNAi^* animals) induced a pupariation delay of 4 days, compared to the 7 days observed in *Tc-torso^RNAi^* larvae (Fig 2A). Moreover, the absence of *Tc-Ptth* did not affect the weight of the resulting pupae when compared to *Control* pupae (Fig 2D and E). Taken together, the differences between *Tc-torso^RNAi^* and *Tc-Ptth^RNAi^* animals suggest that the activation of Torso signaling in *Tribolium* might not rely only on *Tc-Ptth* but also on additional ligands.

**Figure 2.**
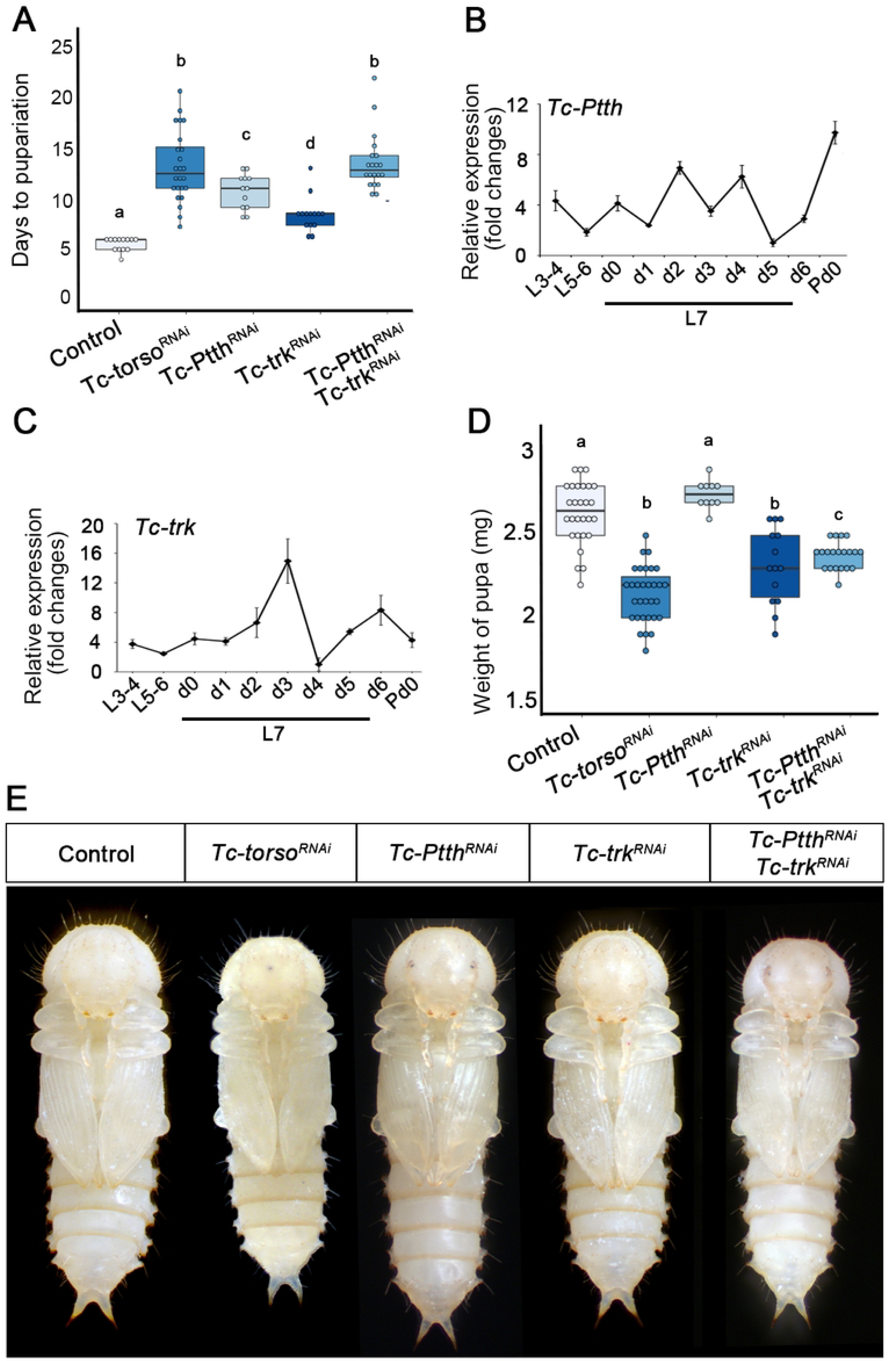
Activation of Tc-Torso by Tc-Ptth and Tc-Trk during the final larval stage of *Tribolium*. (A) Developmental duration of the last larval stage (L7) in newly molted L6 larvae injected with *dsMock* (Control) (n=20), *dsTc-torso* (n=30), *dsTc-ptth* (n=30), *dsTc-trk* (n=23), and *dsTc-ptth* + *dsTc-trk* (n=23). (B) Temporal changes in *Tc-ptth* mRNA and *Tc-trk* mRNA levels measured by qRT-PCR during larval development, from L3-4 to the final (L7) instar larvae. Transcript abundance values are normalized against the Tc-Rpl32 transcript. Error bars indicate the standard error of the mean (SEM) (n=5). (D) Body weight of the last larval stage (L7) of *Control* (n=20), *Tc-torso^RNAi^* (n=32), *Tc-Ptth^RNAi^* (n=31), *Tc-trk^RNAi^* (n=25), and *Tc-Ptth^RNAi^* + *dsTc-trk^RNAi^* (n=21) larvae. All weights were measured on day 5 of the L7 instar. (E) Ventral view of a *Control* and *Tc-torso^RNAi^*, *Tc-Ptth^RNAi^*, *Tc-trk^RNAi^*, and *Tc-Ptth^RNAi^* + *Tc-trk^RNAi^* pupa. Scale bar represents 0.5 mm. In A and D, boxplots are used to represent the data, with black lines indicating medians, colored boxes showing the IQR, and bars indicating the upper and lower values. Values > ±1.5 x the IQR outside the box are considered outliers. Different letters represent groups with significant differences based on an ANOVA test (Tukey, p < 0.001).

In *Drosophila*, the Torso pathway is also required for the formation of the most anterior and posterior regions of the embryo. During this process, Torso receptor is activated by the ligand Trk, which is synthetized in the early embryo [14,25–27]. Similarly, generation of abdominal segments in *Tribolium* requires the Tc-Trk-dependent activation of Torso signaling in the posterior region of the early embryo [26]. Although *Drosophila trk* is not expressed during larval development, ectopic expression of Trk induced a mild advance on pupariation [26], indicating that Trk is able to activate Torso signaling in the larval PG of *Drosophila*. Importantly, we were able to detect expression of *Tc-trk* in the last instar larvae of *Tribolium*, presenting a similar pattern than *Tc-torso*, with a peak of expression at day 3 (Fig 2C). Next, we injected dsRNA of *Tc-trunk* (*Tc-trk^RNAi^* animals) in L6 larvae to analyse its functional relevance during the last larval instar. In contrast to *Ptth*-depleted animals, *Tc-trk^RNAi^* individuals exhibited 2 days of puparium delay (Fig 2A). Importantly, however, the resulting *Tc-trk^RNAi^* pupae were as small as the *Tc-torso^RNAi^* pupae (Fig 2D and E). These results suggest that the activation of Tc-Torso during the last stage of larval development depends on both Tc-Trunk and Tc-Ptth. To confirm this possibility, we knocked-down both ligands simultaneously (*Tc-Ptth^RNAi^* + *Tc-trk^RNAi^* animals). As expected, depletion of *Tc-trk* and *Tc-Ptth* phenocopied the absence of *Tc-torso*, as *Tc-trk^RNAi^* + *Tc-ptth^RNAi^* animals presented 7 days of delay and smaller pupae (Fig 2A, D and E). Altogether, these results show that both ligands, Tc-Trk and Tc-Ptth, act as Tc-Torso ligands to activate Torso signaling during the last larval stage in order to trigger a timely metamorphic transition.

### *Tc-torso* expression is associated with the Threshold Size checkpoint

Our results above strongly suggest that Tc-Torso function is specifically of the last larval instar of *Tribolium*. It is at this stage that *Tribolium* larvae pass through a critical size-assessment checkpoint, the TS, which sets in motion the endocrine and genetic changes that trigger the metamorphic transition at the ensuing molt. The TS checkpoint in

*Tribolium* is reached during the first 24 h after molting to the L7 stage and is associated with the stage-specific down-regulation of *Tc-Kr-h1* and up-regulation of *Tc-Br-C* and *Tc-E93* [33]. Since *Tc-torso* is strongly upregulated during the first 24 h of L7, we wondered whether this increase is also associated to the TS checkpoint. To address this issue, we starved L7 larvae before reaching the TS and measured mRNA levels of *Tc-torso* 72 h later. As Fig 3A shows, *Tc-torso* levels did not increase in larvae starved before the TS, confirming that the stage-specific up-regulation of *Tc-torso* is associated to larvae reaching the TS checkpoint.

**Figure 3.**
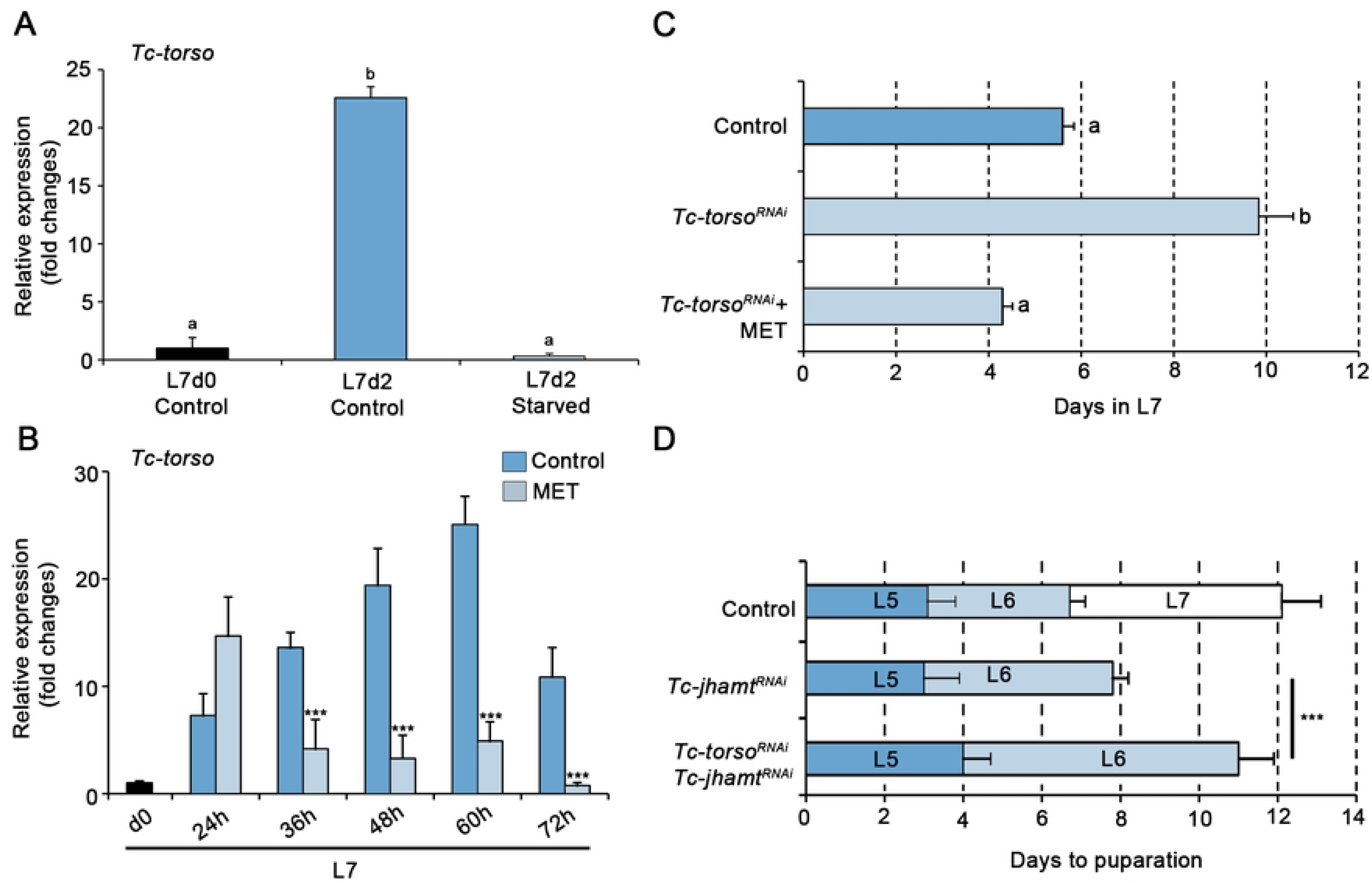
Juvenile Hormone (JH) regulation of *Tc-Torso* expression in *Tribolium*. (A) Temporal changes in *Tc-torso* mRNA levels measured by qRT-PCR in *Control* and starved animals at the indicated stages. Transcript abundance values were normalized against the *Tc-RpL32* transcript. Error bars indicate the standard error of the mean (SEM) (n=5). Asterisks indicate statistically significant differences at ***p ≤ 0.001 (t-test). (B) *Tc-torso* mRNA levels measured by qRT-PCR during the L7 larval instar of *Control* and methoprene-treated animals. Note the dramatic decrease in *Tc-torso* transcription upon methoprene application. Transcript abundance values were normalized against the *Tc-RpL32* transcript. Error bars indicate the SEM (n=5). Asterisks indicate statistically significant differences at ***p ≤ 0.001 (t-test). (C) Developmental duration of the last larval stage (L7) in newly molted L6 larvae injected with *dsMock* (*Control*) (n=20), *dsTc-torso* (n=23), and *dsTc-torso* + methoprene (n=23). The developmental delay induced by depletion of *Tc-torso* is abolished by the application of methoprene. (D) Developmental progression of newly molted L4 larvae after injection with either *dsMock* (*Control*) (n=15), *dsTc-jhamt* (n=25) or *dsTc-torso* + *dsTc-jhamt* (n=21). The bars represent the mean ± standard deviation (SD) for each developmental stage observed after the double-stranded RNA (dsRNA) injection. Error bars indicate the SEM (n=5). Asterisks indicate statistically significant differences at ***p ≤ 0.001 (t-test). Error bars in (A) and (C) indicate the SEM (n=8), and different letters represent groups with significant differences according to an ANOVA test (Tukey, p < 0.001).

Since attainment of the TS is also linked with the decline of JH levels [33], we next wondered if low levels of this hormone are required for the proper expression of *Tc-torso*. To study this, we treated L7 larvae with the JH-mimic methoprene at the TS to maintain high levels of this hormone throughout the larval stage. Under this condition, *Tc-torso* was properly up-regulated just after the TS but its expression significantly decreased thereafter when compared to the progressive increase observed in *Control* larvae (Fig 3B). Consequently, changing the identity of the last larval stage by the application of methoprene, reverted the delay induced by *Tc-torso^RNAi^* and induced the molting to a supernumerary L8 larva (Fig 3B). On the contrary, premature metamorphosis induced by depleting the rate limiting JH biosynthesis enzyme JH acid methyltransferase-3 (Tc-Jhamt) in newly emerged antepenultimate L4 instar larvae (*Tc-jhamt^RNAi^* animals) was delayed by *Tc-torso* depletion. The majority of *Tc-jhamt^RNAi^* larvae molted to normal L5 larvae, and then to L6 underwent precocious metamorphosis after 5 days (Fig 3D). Interestingly, the time to pupation increased significantly when *Tc-jhamt* and *Tc-torso* were depleted simultaneously (*Tc-jhamt^RNAi^* + *Tc-torso^RNAi^* animals) (Fig 3D), thus confirming that *Tc-torso* function does not depend on the number of larval stages the larva has been trough but on whether the animal is in its last larval stage. Altogether, these results indicate that (1) the early upregulation of *Tc-torso* in L7 requires the larvae reaching the TS checkpoint; and (2) the ensuing increase in *Tc-torso* expression must occur in the presence of very low levels of JH.

### TGFß/Activin signaling pathway regulates expression by repressing JH synthesis

Since the up-regulation of *Tc-torso* correlates with the decline in JH levels triggered by the TS checkpoint, we next wanted to study the relation between both processes. In this regard, it has been recently shown that the TGFß/Activin signaling pathway is responsible for the decline of JH levels in a number of insect species [30–32]. We, therefore, wanted to ascertain whether the TGFß/Activin pathway regulates the expression of *Tc-torso* in the last larval instar. We first examined the expression of the Activin-like ligand Myoglianin (Tc-Myo) and its Type-I receptor Baboon (Tc-Babo) during the penultimate and last larval stages. Whereas *Tc-babo* mRNA levels persisted without major fluctuations through the last two larval instars, those of *Tc-myo* were more dynamic with a remarkable increase during the first two days and then oscillating throughout the rest of the instar (Fig 4A and B). We then depleted *Tc-babo* and *Tc-myo* during the last larval instar by injecting the corresponding dsRNAs (*Tc-babo^RNAi^* and *Tc-myo^RNAi^* animals). Remarkably, we found that in contrast to the normal upregulation of *Tc-torso* observed in *Control* larvae two days after the TS checkpoint, the levels of *Tc-torso* in *Tc-babo^RNAi^* and *Tc-myo^RNAi^* larvae were dramatically reduced, being even lower than the levels observed in newly molted L7 control animals (Fig 4C). In addition, *Tc-babo^RNAi^* and *Tc-myo^RNAi^* larvae presented persistently elevated levels of *TcKr-h1*, rather than the low levels observed in *Control* larvae (Fig 4D), which is indicative of sustained high levels of JH in these animals. Consistently, *Tc-E93* and *Tc-Br-C* expression did not increase in *Tc-babo^RNAi^* and *Tc-myo^RNAi^* larvae after reaching the TS (Fig 4E and F). Altogether these results show that the activity of TGFß/Activin signaling pathway at the onset of L7 is responsible for the decline in JH levels that triggers the changes in the expression of the temporal factors that control the metamorphic transition, including the sharp up-regulation of *Tc-torso*.

**Figure 4.**
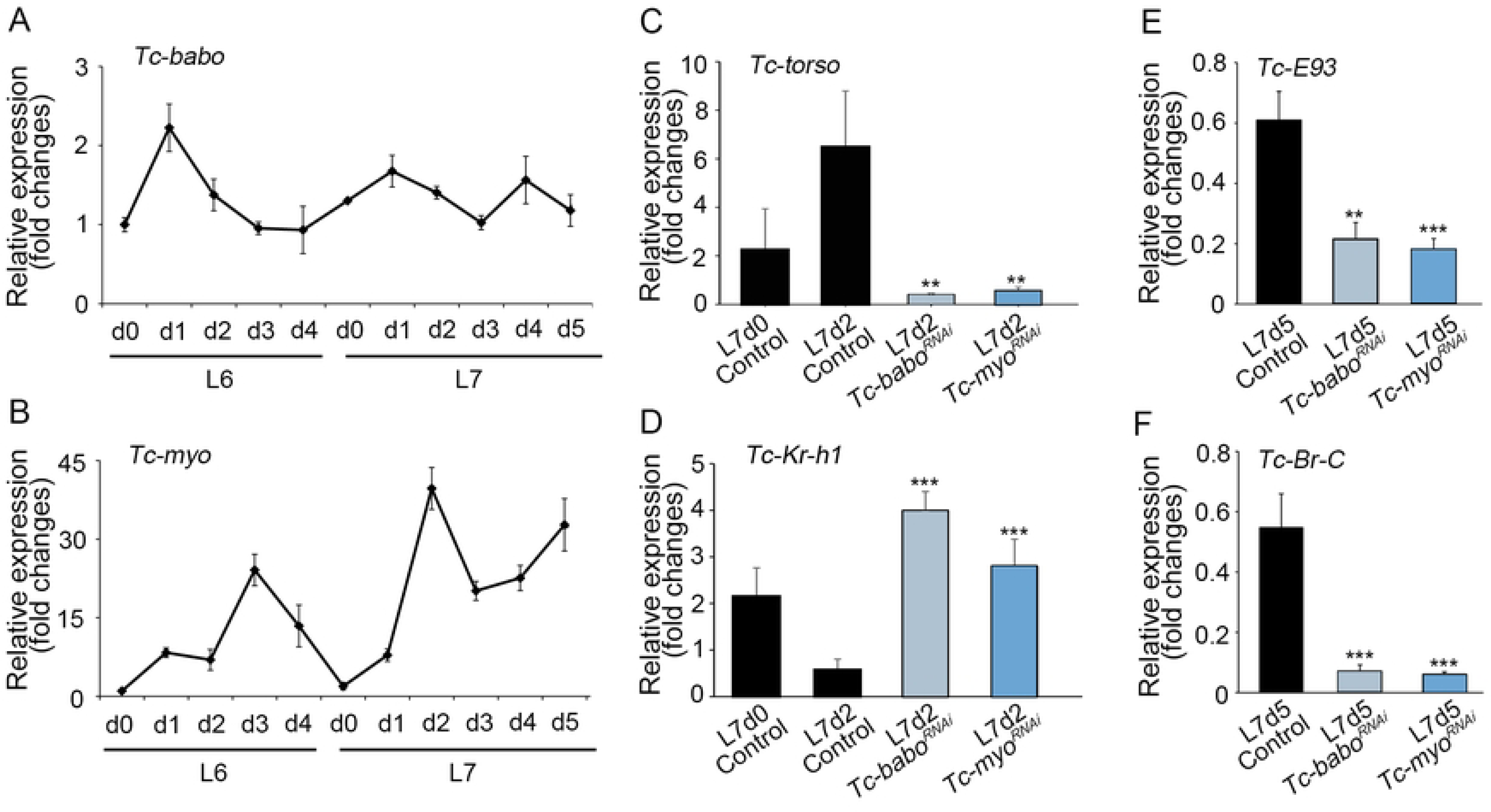
TGFß/Activin signaling pathway activates *Tc-torso* expresión through the repression of JH synthesis. (A-B) Temporal changes in *Tc-babo* (A) and *Tc-myo* (B) mRNA levels measured by qRT-PCR in penultimate (L6) and ultimate (L7) instar larvae. Transcript abundance values were normalized against the *Tc-Rpl32* transcript. Error bars indicate the standard error of the mean (SEM) (n=5). (C-D) Temporal changes in transcript levels of *Tc-torso* (C), *Tc-Kr-h1* (D), *Tc-E93* (E), and *Tc-Br-C* (F) measured by qRT-PCR at the indicated time points of L7 *Control*, *Tc-babo^RNAi^*, and *Tc-myo^RNAi^* larvae. Transcript abundance values were normalized against the *Tc-Rpl32* transcript. Average values of three independent datasets are shown with standard errors (n=6). Asterisks indicate statistically significant differences at **p < 0.01 and ***p < 0.001 (t-test).

### The TGFß/Activin signaling pathway facilitates the metamorphic transition by regulating ecdysone levels

The results above indicate that *Tc-babo^RNAi^* and *Tc-myo^RNAi^* larvae do not initiate the molecular events that characterize the nature of the last larval instar, which involve the decline in *Tc-Kr-h1* expression with the concomitant upregulation of *Tc-E93*, *Tc-Br-C* and *Tc-torso*. It is expected, therefore, that *Tc-babo^RNAi^* and *Tc-myo^RNAi^* larvae would not initiate the larva-pupa transformation at the ensuing molt. Consistent with this and in agreement with a previous study [32], *Tc-myo^RNAi^* larvae failed to pupate and instead repeated successive larval molts to ultimately reach L10 instar larva (when we stop analyzing the knockdown animals). Remarkably, the intermolt period was significantly increased in these larvae (Fig 5A and B), and the weight of the supernumerary *Tc-myo^RNAi^* larvae remained above the TS (Fig 5C), indicating that metamorphosis cannot be triggered in the absence of active TGFß/Activin signaling. On the other hand, the phenotype of *TcBabo^RNAi^* larvae was even more dramatic as they never molted again and remained as L7 larvae an average of 65 days before dying (Fig 5D). The phenotypes presented by *Tc-babo^RNAi^* and *Tc-myo^RNAi^* larvae suggest that TGFß/Activin signaling pathway might play a role in controlling the synthesis of ecdysone, as previously reported in *Drosophila*, in addition to the regulation of JH [29]. Supporting this possibility, the levels of the Halloween gene *Tc-phm* and the ecdysone-dependent gene *Tc-HR3*, used as proxies for 20E levels, were significantly downregulated in *Tc-babo^RNAi^* and *Tc-myo^RNAi^* larvae when compared to *Control* animals (Fig 5E and E), which demonstrate that TGFß/Activin signaling is required for ecdysone synthesis in *Tribolium*. Colectively, our findings reveal that TGFß/Activin signaling exerts a dual role in the control of metamorphic timing in *Tribolium*: firstly, it is responsible for the decline in JH levels at the TS checkpoint, triggering the genetic switch that initiate the metamorphic transition, including the up-regulation of *Tc-torso*; secondly, it controls the production of ecdysone, a critical factor required to elicit the corresponding developmental transition.

**Figure 5.**
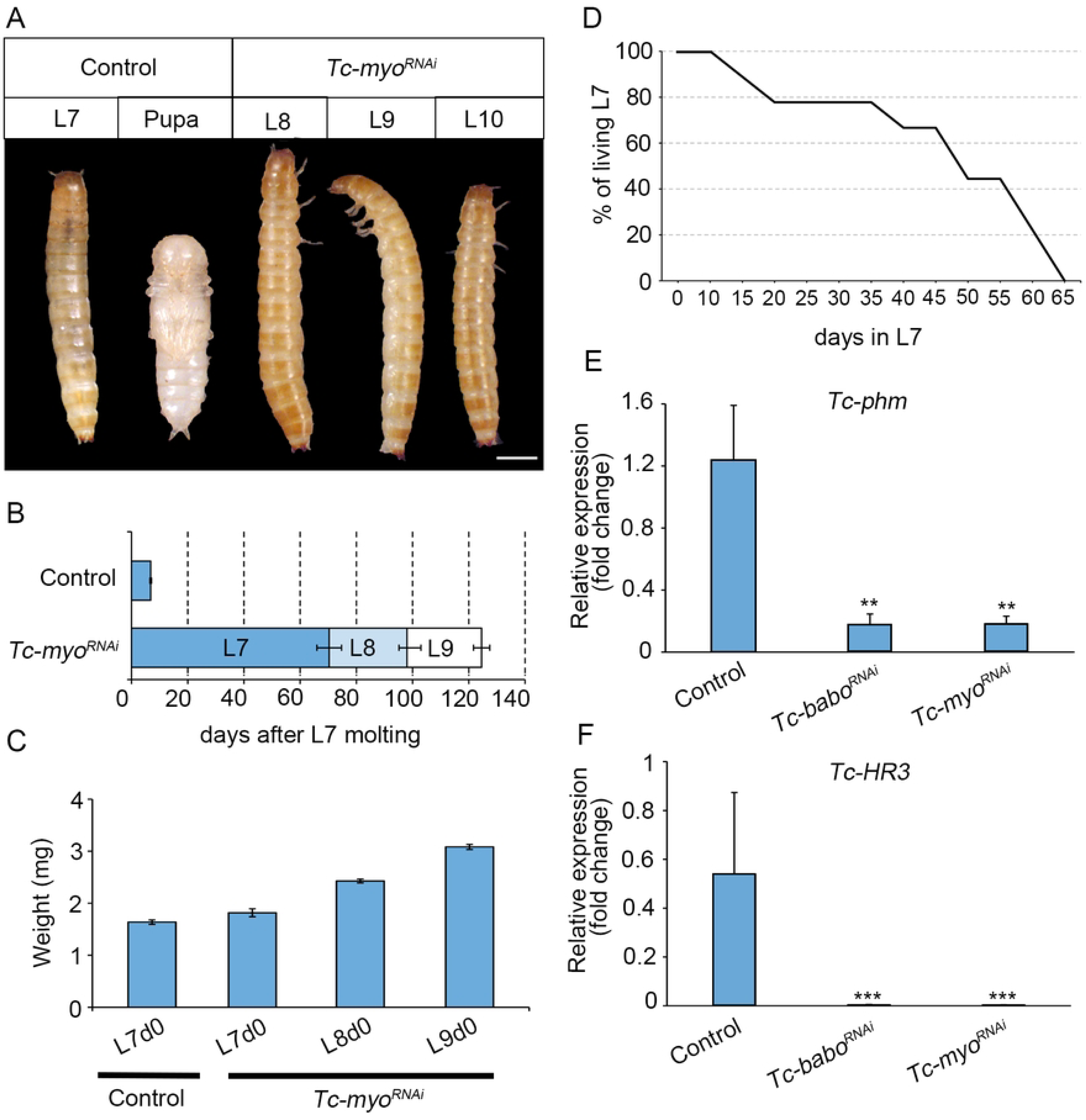
Impairment of metamorphosis upon inactivation of TGFß/Activin signaling. (A) Dorsal and ventral view of a Control L7 larva and pupa, and dorsal views of supernumerary L8, L9, and L10 *Tc-torso^RNAi^* animals. Scale bar, 0.5 mm. (B) Temporal progression of L6 larvae injected with *dsMock* (Control) (n=30) or with *dsTc-myo* (n = 81) and left until the ensuing molts. Each bar indicates the periods (mean ± SD) for each developmental stage after the molt into the L7 larval stage. (C) Growth in body weight of *Control* and *Tc-myo* animals. All weights are measured on day 0 of each instar. Bars indicate the mean ± SD. (D) Number of *Tc-babo* depleted larvae alive recorded for each day after the molt into the L7 larval stage. (E) Transcript levels of *Tc-phm*, and *Tc-HR3* measured by qRT-PCR in 2-day-old L7 Control, *Tc-babo^RNAi^* and *Tc-myo^RNAi^* larvae. Transcript abundance values were normalized against the *Tc-Rpl32* transcript. Average values of three independent datasets are shown with standard errors (n=6). Asterisks indicate statistically significant differences at **p < 0.01 and ***p < 0.001 (t-test).

## Discussion

Here, we show that the Ptth-Trunk/Torso signaling pathway plays a crucial role in the transition from juvenile to adult stages in *Tribolium*. Our data reveal that both Ptth and Trk ligands activate Tc-Torso signaling during the final larval stage of the beetle. However, whereas Ptth promotes ecdysone biosynthesis, Trk promotes organ growth. Interestingly, we found that the expression of *Tc-torso* depends on the TGFβ/Activin signaling-dependent decay of JH at the TS checkpoint. In addition, we also found that TGFβ/Activin signaling contributes to ecdysone biosynthesis. These findings improve our understanding of the molecular mechanisms underlying the hormonal control of metamorphic transitions and provide insights into the evolution of Torso signaling in insects.

### Postembryonic Torso signaling is activated by PTTH and Trk in *Tribolium*

Contrary to previously studied insects, our results demonstrated that post-embryonic Torso signaling in *Tribolium* is activated by both Ptth and Trk. To date, Ptth was the only ligand known to activate Torso during the post-embryonic stages in insects [14,35], whereas Trk was described as a Torso ligand exclusively involved in the generation of the terminal structures during the early stages of embryogenesis of *Drosophila* and *Tribolium* [25,26]. Despite the temporal specificity of the activation of the signaling by each ligand, it is important to note that both ligands seem to be functionally interchangeable, as ectopic expression of Ptth is able to activate Torso signaling in the early embryo of *Drosophila* [14], and overexpression of a processed form of Trk is able to induce precocious metamorphosis when overexpressed in the PG of the fly [26]. This indicates a possible common origin of both ligands, which have acquired specific enhancers to activate the pathway in different tissues. The fact that Ptth appears to evolve from the original ligand Trk in the common ancestor of Hemiptera and Holometabola supports this idea [36].

Although we found that Tc-Trk and Tc-Ptth activates Torso signaling during the metamorphic transition in *Tribolium*, both ligands present common and non-overlapping functions. Whereas Tc-Ptth is involved in the regulation of ecdysone production and the timely induction of the metamorphic transition, Trk regulates the systemic growth during the last larval stage. Interestingly this data is consistent with results obtained in *Drosophila* and *Bombyx,* where inactivation of Torso specifically in the PG delays the onset of pupariation, extending the larval growth period and increasing the final pupal size [13,14,37], whereas depletion of *Torso* specifically in the fat body produced smaller pupae with no effect on developmental timing [38]. Although the ligand responsible for Torso activation in the *Drosophila* fat body has not been identified, these results raise the possibility that Trk might activate Torso in the fat body of *Tribolium* to regulate growth rate. Generation of new genetic tools to deplete gene expression in a tissue-specific manner would be required to solve this question in *Tribolium*.

In *Drosophila* and in *Bombyx*, *torso* mutants produced larger adults due to a prolonged larval development that allows for extra growth [13,14,37]. However, our study shows that depletion of *Tc-Torso* in *Tribolium* reduces final pupal size even when the larval growth period is extended. This finding suggests that the activation of Tc-Torso by Tc-Trk in the fat body may play a crucial role in increasing larval body mass. Considering that Trk and Torso are the most ancient molecules of the signaling pathway, as indicated by their presence in chelicerates, the most basal group of arthropods [36], it is tempting to speculate that systemic growth regulation is likely the ancestral function of the pathway. The fact that co-option of Torso signaling in early embryogenesis as well as for ecdysone biosynthesis during post-embryonic development seems to be a relative evolutionary novelty in insect evolution support this idea [27,36]. Studying the function of Trk-Torso in hemipteran species and Trk-like molecules in chelicerates could shed light on this possibility.

### Up-regulation of *Tc-torso* depends on the TS checkpoint

Our expression analysis of *Tc-torso* in *Tribolium* reveals a peak of transcription at the TS checkpoint. This peak coincides with the attainment of the TS at the beginning of the final larval stage, accompanied by a decline in JH levels. The decrease in JH levels enables the up-regulation of Tc-E93 and Tc-Br-C, initiating the metamorphic transition. Interestingly, during the last larval stage an important increase of ecdysone production is required to induce the quiescent stage and the subsequent pupa formation [39]. In this sense, the decrease of JH production has been related to the upregulation of the Halloween genes, responsible for ecdysone biosynthesis. Thus, in *Drosophila*, inactivation of JH signalling in the PG triggers premature metamorphosis by de-repression of Halloween genes [40]. Such effect of JH on ecdysone biosynthesis might depend on the precocious up-regulation of *Torso* expression since ecdysone production relays in part on the activation of Torso signalling. Indeed, our results show that application of JH in the last larval stage of *Tribolium* impairs *Tc-torso* up-regulation. This effect has been also observed in *Bombyx* where the application of a JH analog not only downregulates *Bm-torso* but also several Halloween genes [41]. These observations, therefore supports the idea that JH mainly represses ecdysone production by regulating *torso* expression.

### TGFβ/Activin signaling regulates JH biosynthesis

If the induction of *torso* expression depends on the disappearance of JH in the last larval stage, then what regulates such decline? Interestingly, in agreement with a previous report [32] we found that inhibition of JH synthesis depends on the activation of TGFβ/Activin signaling, as depletion of either its ligand *Tc-myo* or the main receptor *Tc-babo* blocks the metamorphic transition. Consistently, blocking the TGFβ/Activin signaling leads to a sustained high levels of the JH-dependent factor *Tc-Kr-h1*, indicating that JH levels do not decline under these circumstances. Similar results have been reported in other insects, suggesting a conserved role of TGFβ/Activin signaling in JH regulation. Thus, in *H. vigintioctopunctata*, *G. bimaculatus* and *B. germanica* inactivation of TGFβ/Activin signaling blocks JH production by directly downregulation of the JH biosynthetic enzyme gene, *jhamt* [31,32,42]. The fact that JH regulation by TGFβ/Activin signaling occurs specifically during the last larval stage suggests a potential relationship between body mass and the activation of TGFβ/Activin signaling. Interestingly, in the lepidopteran *Manduca,* increasing levels of *Ms-myo*, produced by the muscle, have been associated with the initiation of metamorphosis [32]. This data strongly suggests that muscle growth during larval development induces a gradual increasing of *myo* expression that activates TGFβ/Activin signaling upon reaching a certain threshold level. Consequently, this activation leads to a decrease of JH production, allowing for the up-regulation of *Torso* and the subsequent raise of ecdysone levels. Such complex regulation might explain why Torso is only required during the last larval stage.

### TGFβ/Activin signaling as a PG cell survival factor

In addition to its role in regulating JH, TGFβ/Activin signaling also plays a role in ecdysone production in *Tribolium*, as revealed by *Tc-myo* knockdown. In agreement with a previous report [32], we found that depletion of *Tc-myo* in *Tribolium*, increased dramatically the inter-molting period and even induced a developmental arrest at L7 when the TGFβ/Activin receptor *Tc-babo* was knockdown. These results are reminiscent of the developmental arrest observed in *Drosophila* larvae when TGFβ/Activin signaling is inactivated in the PG [29]. As in *Drosophila*, arrested *Tc-babo-*depleted *Tribolium* larvae presented very low levels of *Tc-torso* expression with the consequent failure to induce the large rise in ecdysteroid titer that triggers metamorphosis (Fig 4C). Under these conditions, we anticipated the presence of supernumerary larvae in the subsequent molt due to the failure of JH decay. However, we found that inactivation of TGFβ/Activin signaling in *Tribolium* not only halts metamorphosis but also prevents larva molting, as depletion of *Tc-babo* leads to developmental arrest at L7 for more than 60 days. Considering that the effect of the injected dsRNAs is transient [43], the most likely explanation for the arrested *Tc-babo*-depleted larvae is that TGFβ/Activin might be required for PG viability during the last larval stage. This hypothesis is supported by the fact that inactivation of TGFβ/Activin signaling in adult *Drosophila* muscles reduces lifespan by modulating protein homeostasis in the tissue [44]. Likewise, reduction of TGFβ/Activin signaling in the cardiac muscle of *Drosophila* increases the cardiac autophagic activity [45]. Since autophagy is the main mechanism for larval tissue degradation during metamorphosis [46], it is tempting to speculate that TGFβ/Activin activation in the PG suppresses autophagy during the last larval stage of *Tribolium*. The characterization of the PG in *Tribolium* and specific analysis of the effects on TGFβ/Activin signaling in this tissue will provide further insights into this question.

In summary, our study reveals that the activation of Tc-Torso by two ligands, Tc-Ptth and Tc-Trk, during the last larval stage of *Tribolium* regulates the ecdysone-dependent timely transition to the metamorphic period and controls growth during this stage, thereby determining the final size of the insect. Furthermore, we showed *Tc-torso* up-regulation depends on larvae attaining the TS checkpoint at the onset of the last larval instar and that its expression depends on the decay of JH induced by the TGFβ/Activin signaling pathway. Our findings, therefore contribute to the understanding of how the coordination of different signaling pathways regulates the endocrine systems that control developmental growth and the timing of maturation in insects.

## Materials and methods

### Tribolium castaneum

The enhancer-trap line pu11 of Tribolium (obtained from Y. Tomoyasu, Miami University, Oxford, OH) was reared on an organic wheat flour diet supplemented with 5% nutritional yeast and maintained at a constant temperature of 29 °C in complete darkness.

### Quantitative Real-Time Reverse Transcriptase Polymerase Chain Reaction (qRT-PCR)

Total RNA from individual *Tribolium* larvae was extracted using the GenEluteTM Mammalian Total RNA kit (Sigma). cDNA synthesis was performed following the previously described methods [47,48]. For quantitative real-time PCR (qPCR), Power SYBR Green PCR Mastermix (Applied Biosystems) was used to determine relative transcript levels. To standardize the qPCR inputs, a master mix containing Power SYBR Green PCR Mastermix and forward and reverse primers was prepared, with each primer at a final concentration of 100 µM. The qPCR experiments were conducted with an equal quantity of tissue equivalent input for all treatments, and each sample was run in duplicate using 2 µl of cDNA per reaction. As a reference, the same cDNAs were subjected to qRT-PCR using a primer pair specific for *Tribolium* Ribosomal *Tc-Rpl32*. All samples were analyzed on the iCycler iQReal Time PCR Detection System (Bio-Rad).

### Primer sequences used for qPCR for *Tribolium*

*Tc-torso-F*: 5’-TTGACGAGGAGAAGCTTCCAGAGT-3’

*Tc-torso-R*: 5’-TGCAAATTGTTGCTGCATGTTGGT-3’

*Tc-Ptth-F*: 5’-TCGTGTGGGATCGAATTTCGCGTTC-3’

*Tc-Ptth-R*: 5’-GTCTTTCTGTTTCAAAACGCGGAC-3’

*Tc-trk-F*: 5’-TATCGAGCAACTCGACGAT-3’

*Tc-trk-R*: 5’-TCGATTTGCAATGCCACTGTT-3’

*Tc-myo-F*: 5’-CAAGAAGTGCTCACCTTTGC-3’

*Tc-myo-R*: 5’-CCTTCATGTACACGTACAG-3’

*Tc-babo-F*: 5’-ATGGTGCATGGCTTCTGGTT-3’

*Tc-babo-R*: 5’-AGTAAGTCGATGTAAAGCAGTA-3’

*Tc-E75-F*: 5′-CGGTCCTCAATGGAAGAAAA*-*3′

*Tc-E75-R*: 5′*-*TGTGTGGTTTGTAGGCTTCG-3′

*Tc-phm-F*: 5′-TGAACAAATCGCAATGGTGCCATA*-*3′

*Tc-phm-R*: 5′*-*TCATGGTACCTGGTGGTGGAACCTTAT-3′

*Tc-Kr-h1-F*: 5′-AATCCTCCTGCTCATCCAGCACTA-3′

*Tc-Kr-h1-R*: 5′-CAGGATTCGAACTAGGAGGTGTTA-3′

*Tc-E93-F*: 5′-CTCTCGAAAACTCGGTTCTAAACA-3′

*Tc-E93-R*: 5′-TTTGGGTTTGGGTGCTGCCGAATT-3′

*Tc-Br-C-F*: 5′-TCGTTTCTCAAGACGGCTGAAGTG*-*3′

*Tc-Br-C-R*: 5′*-*CTCCACTAACTTCTCGGTGAAGCT-3′

*Tc-Rpl32-F*: 5′-CAGGCACCAGTCTGACCGTTATG*-*3′

*Tc-Rpl32-R*: 5′*-*CATGTGCTTCGTTTTGGCATTGGA-3′

### Larva RNAi injection

*Tc-torso* dsRNA (IB_04720), *Tc-Ptth* dsRNA (IB_09326), *Tc-Trk* dsRNA (IB_06187), Tc-myo dsRNA (IB_05899), *Tc-jhamt* dsRNA (IB_04499), and *Tc-babo* dsRNA (IB_03525) were synthesized by Eupheria Biotech Company. The control dsRNA used a non-coding sequence from the pSTBlue-1 vector (dsMock). For larval injections, a concentration of 1 µg/µl dsRNA was administered to penultimate instar and antepenultimate instar larvae. In cases of co-injection of two dsRNAs, equal volumes of each dsRNA solution were mixed and applied in a single injection.

### Nutritional Experiments

Tribolium pupae-larvae were reared on a normal diet consisting of organic wheat flour containing 5% nutritional yeast. For the starvation experiment, newly molted L7 larvae, raised on the normal diet, were transferred to a new plate without any food. After 2 days, the larvae were collected for *Tc-torso* expression measurement using RT-qPCR.

### Treatment with Methoprene

To conduct the juvenile hormone mimic treatment, white L7 newly molted larvae of Tribolium were topically treated on their dorsal side with 1μg of isopropyl (E,E)-(RS)-11-methoxy-3,7,11-trimethyldodeca-2,4-dienoate per specimen in 1 μL of acetone. The control group received the same volume of solvent. At the desired stage, the larvae were subjected to mRNA expression analysis.

### Microscopy analysis

All pictures were obtained with AxioImager.Z1 (ApoTome 213 System, Zeiss) microscope, and images were subsequently processed using Fuji and Adobe photoshop.

## Acknowledgements

This project is supported by grants PGC2018-098427-B-I00 and PID2021-125661NB-100 to D.M., and X.F-M. funded by MCIN/AEI/10.13039/501100011033 and by grant 2017-SGR 1030 and 2021-SGR 0417 to D.M. and X.F-M funded by the Secretaria d’Universitats i Recerca del Departament d’Economia i Coneixement de la Generalitat de Catalunya. The research has also benefited from ERDF “A way of making Europe” to D.M. and X.F-M. S.C. is a recipient of a Juan de la Cierva FJC2019-041549-I contract from the MCIN.

